# The Importance of Anti-Correlations in Graph Theory Based Classification of Autism Spectrum Disorder

**DOI:** 10.1101/557512

**Authors:** Amirali Kazeminejad, Roberto C Sotero

**Affiliations:** Department of Biomedical Engineering, University of Calgary, Calgary, Alberta, Canada; Hotchkiss Brain Institute, University of Calgary, Calgary, Alberta Canada; Department of Radiology, University of Calgary, Calgary, Alberta, Canada

## Abstract

With the release of the multi-site Autism Brain Imaging Data Exchange, many researchers have applied machine learning methods to distinguish between healthy subjects and autistic individuals by using features extracted from resting state functional MRI data. An important part of applying machine learning to this problem is how to extract these features. Specifically, whether to include anti-correlations between brain regions as relevant features and how best to define these features. For the second question, the graph theoretical properties of the brain network may provide a reasonable answer. In this study, we investigated the first issue by comparing three different approaches. These included using the unaltered correlation matrix, the absolute value of the correlation matrix or the inverted correlation matrix (negative of the original) as the starting point for extracting relevant features using graph theory. We then trained a multi-layer perceptron in a leave-one-site out manner in which the data from a single site was left out as testing data and the model was trained on the data from the other sites. Our results show that on average, using graph features extracted from the anti-correlation-based matrix led to the highest accuracy and AUC scores. We also show that adding the PCA transformation of the original correlation matrix to the feature space leads to an increase in accuracy. This suggests that anti-correlations should not simply be discarded as they may include useful information that would aid the classification task.

## 1 Introduction

Autism Spectrum Disorder (ASD) is a neurodevelopmental condition that is growing in prevalence in recent years [1]. While it is usually diagnosed by carefully monitoring a child’s behavioral development [2], recent studies have shown that brain imaging can also be used to aid in that diagnosis by identifying underlying differences between the ASD and Healthy Control (HC) Brain[3], [4].

Functional Magnetic Resonance Imaging (fMRI) is one of the most widely used tools for such studies due to its high spatial resolution. It monitors the changes in the Blood Oxygen Level Dependent (BOLD) signal which indirectly measures the neuronal activity [5]. This allows researchers to examine the human brain at a network level and understand how each brain region is connected to the rest of the brain by analyzing the BOLD activation patterns. A variant of the technique call Resting State fMRI (rs-fMRI) has been widely used to examine brain networks while subjects are at rest [6], [7].Graph theory is one of the more novel methods being used for the network-level analysis of the brain. It provides a mathematical framework for quantifying network characteristics and quantitively analyze the differences between different brain networks [8], [9]. Machine learning is another relatively new technique that is being applied to rs-fMRI data to extract insights such as important biomarkers as well as to develop novel algorithms with the hope of automatically diagnosing brain disorders from medical imaging data. [10], [11]. The network characteristics that is provided by graph theory has been used as the input to machine learning algorithms to identify diseases such as Alzheimer’s [12], Parkinson’s [12], [13], and Autism [14].

One of the variables that can affect the results of the analysis is how the graph construction step was performed. After the initial preprocessing of the rs-fMRI data, a correlation matrix is created by calculating the correlation of each brain region with the other regions. The correlation matrix is then transformed into a sparse binary matrix representing the existence connections between different regions. This transformation is usually performed by a thresholding step in which only the two strongest correlations are kept [8], [9]. However, this step ignores the anti-correlations which may be biologically relevant [15]. This problem will be even more severe if the preprocessing pipeline includes the regression of the global mean signal (GSR) from the time-series, known to result in the removal of motion, cardiac and respiratory signals, which may result in the presence of more anticorrelations in the correlation matrix [16]. The use of GSR is a controversial matter in the field. Although there is evidence suggesting that the anticorrelations introduced through GSR have no biological basis [17], a recent study has shown using GSR in the preprocessing pipeline for rs-fMRI leads to better prediction accuracies of behavioral measures [18].

There has been evidence of network-level changes in the ASD brain compared to a HC brain[4], [19], [20]. Therefore, using graph theory to extract features for classification of ASD is likely to provide good results. However, this approach is affected by how the graph theory analysis is formulated. This claim was previously examined by training a SVM classifier on graph theoretical features to classify between ASD and HC [14]. However, the methodology of that paper was limited to using only the positive correlations in order to construct the brain graph, potentially ignoring some informative connections.

The release of the Autism Brain Imaging Data Exchange I (ABIDE I) dataset[21] allowed researchers to examine ASD in large sample sizes. Nielsen et al. [22] conducted one of the earliest classification studies on ABIDE and were able to achieve an accuracy of 60% over all samples. More recently, Heinsfeld et al [23] were able to achieve an accuracy of 70%, evaluated by 10 fold cross-validation, on the entire dataset by utilizing neural networks and transfer learning. They Also reported the leave-one-site out performance of their model averaging at 65% accuracy. Plitt et al [24] achieved a higher accuracy, 69.71% when using only 178 subjects from the ABIDE I dataset.

A recent study by Hallquist et al. [25], examining 106 papers (with only 2 papers focusing on ASD (using graph theoretical measures, shows that 79.2% of graph theoretical studies either did not specify how they handled negative correlations or discarded them. Another 9.4% used absolute values of the correlation matrix. However, this is mostly an arbitrary choice.

In this paper, we study the effect of different approaches to handling anti-correlations for the classification accuracy of a machine learning model on the ABIDE dataset. We trained a regularized deep learning neural network using features extracted from transforming the correlation matrix using three pipelines: The positive correlation pipeline which does not change the matrix, the anti-correlation pipeline which prioritizes negative correlations, and the absolute value pipeline which disregards the sign of the correlations. Our model was evaluated using a leave-one-site out approach. Our results show that on average, the anti-correlation pipeline results in better accuracy and area under curve (AUC) score with a small loss in sensitivity. Furthermore, our model performed on-par with previous state-of the art models [23] when using only graph theoretical features. Interestingly, adding the PCA transformation of the original correlation matrix increased the accuracy, sensitivity, specificity and AUC score for all pipelines.

## 2 Materials & Methods

### 2.1 Data & Preprocessing

This study used a publicly available dataset from the ABIDE Initiative [26]. To ensure that our results were not affected by any custom preprocessing pipeline, we used the preprocessed data provided by ABIDE in the C-PAC [27] pipeline. The preprocessing included the following steps. The Analysis of Functional Neuro Images (AFNI)[28] software was used for removing the skull from the images. The brain was segmented into three tissues using FMRIB Software Library (FSL) [29]. The images were then normalized to the MNI 152 stereotactic space[30][31] using Advanced Normalization Tools (ANTs) [32]. Functional preprocessing included motion and slice-timing correction as well as the normalization of voxel intensity. Nuisance signal regression included 24 parameters for head motion, CompCor[33] with 5 principal components for tissue signal in Cerebrospinal fluid and white matter, linear and quadratic trends for Low-frequency drifts and a global bandpass filter (0.01 to 0.1 Hz). GSR was also applied to remove the global mean from the signals. These images where then co-registered to their anatomical counterpart by FSL. They were then normalized to the MNI 152 space using ANTs. The average voxel activity in each Region of Interest (ROI) of the Craddock 200 atlas [34] was then extracted as the time-series for that region. We also used data from the anatomical automatic labeling(AAL)[35] in order to replicate our results on the Craddock 200 atlas. This resulted in data from 1035 subjects. The demographics of whom are outlined in Table 1.

**Table 1.** Subject demographics. M: Male, F: Female, ASD: Autism spectrum disorder, HC: Healthy control, SD: Standard deviation. Some sites did not include female subjects, these have been identified with a “_”. The term “X” for SD means that there was not enough samples to calculate SD.

|  | Age Mean (SD) |  |  |  | Count |  |  |  |
| --- | --- | --- | --- | --- | --- | --- | --- | --- |
| Group | ASD |  | HC |  | ASD |  | HC |  |
| Sex | F | M | F | M | F | M | F | M |
| Site |  |  |  |  |  |  |  |  |
| **CALTECH** | 24.35<br>(6.91) | 28.27<br>(11.08) | 22.77<br>(4.31) | 29.51<br>(11.83) | 4 | 15 | 4 | 14 |
| **CMU** | 27.00<br>(6.93) | 26.18<br>(5.88) | 26.00<br>(5.29) | 27.10<br>(6.12) | 3 | 11 | 3 | 10 |
| **KKI** | 9.61<br>(1.85) | 10.12<br>(1.38) | 9.49<br>(0.81) | 10.22<br>(1.23) | 4 | 16 | 8 | 20 |
| **LEUVEN** | 14.00<br>(1.41) | 18.18<br>(5.09) | 13.56<br>(0.75) | 19.01<br>(5.05) | 3 | 26 | 5 | 29 |
| **MAX_MUN** | 35.33<br>(8.50) | 24.76<br>(15.27) | 32.00<br>(X) | 24.37<br>(8.81) | 3 | 21 | 1 | 27 |
| **NYU** | 18.62<br>(9.14) | 14.14<br>(6.59) | 15.37<br>(6.45) | 15.76<br>(6.10) | 10 | 65 | 26 | 74 |
| **OHSU** | – | 11.43<br>(2.18) | – | 10.09<br>(1.12) | – | – | – | 14 |
| **OLIN** | 19.00<br>(5.00) | 16.06<br>(3.04) | 17.50<br>(3.54) | 16.54<br>(3.78) | 3 | 16 | 2 | 13 |
| **PITT** | 12.60<br>(0.95) | 20.03<br>(7.38) | 16.07<br>(3.60) | 19.36<br>(6.97) | 4 | 25 | 4 | 23 |
| **SBL** | – | 35.00<br>(10.43) | – | 33.73<br>(6.61) | – | 15 | – | 15 |
| **SDSU** | 12.44<br>(X) | 14.89<br>(1.70) | 13.26<br>(2.69) | 14.58<br>(1.46) | 1 | 13 | 6 | 16 |
| **STANFORD** | 10.02<br>(1.63) | 9.99<br>(1.68) | 9.01<br>(1.01) | 10.19<br>(1.66) | 4 | 15 | 4 | 16 |
| **TRINITY** | – | 16.81<br>(3.17) | – | 17.08<br>(3.77) | – | 22 | – | 25 |
| **UCLA** | 12.46<br>(3.42) | 13.08<br>(2.34) | 12.82<br>(0.89) | 13.03<br>(2.04) | 6 | 48 | 6 | 38 |
| **UM** | 13.53<br>(3.12) | 13.11<br>(2.30) | 14.78<br>(2.82) | 14.82<br>(3.85) | 9 | 57 | 18 | 56 |
| **USM** | – | 23.46<br>(8.33) | – | 21.29<br>(8.35) | – | 46 | – | 25 |
| **YALE** | 12.88<br>(3.00) | 12.70<br>(3.14) | 13.52<br>(2.64) | 12.34<br>(2.79) | 8 | 20 | 8 | 20 |

### 2.2 Network Construction, Graph Extraction and Feature Extraction

To construct the brain network, the timeseries for each atlas ROI where correlated, using Pearson correlation, with the other regions. The strengths of these correlations were used as the strengths of the connection between different ROIs. In graph terms, this represents a fully connected graph with each of the nodes being in the center of the corresponding ROIs and each edge weight is the correlation between the two nodes on the opposite ends of the edge. Three different pipelines were obtained based on selecting only the positive values, only the negative values, or taking the absolute value of the correlation matrix. These graphs were then subjected to a thresholding step in which only the 20-50% strongest connections were set to one and the other edges were discarded. This threshold was incremented in 2% steps. This resulted in sparse binary graphs. This step was done because binary graphs have been shown to have more easily defined null models and are more easily characterizable [8]. Several measures of integration (characteristic path length and e□ciency), segregation (clustering coe□cient and transitivity), centrality (betweenness centrality, eigenvector centrality, participation coe□cient and within module z-score) and resilience(assortativity) [9] were then extracted from the binary brain network graph. These steps were done using the Python libraries brainconn (https://github.com/FIU-Neuro/brainconn) and network (https://networkx.github.io/). This resulted in 3 feature sets of 1404 features for each group. The main body of this paper focuses mainly on only using the graph features in the analysis. In the supplementary material, we report the results of adding the PCA transformation [36] of the unaltered correlation matrix to the graph features.

### 2.3 Leave-One-Site-Out (LOSO) Cross-Validation

The structure of the ABIDE I dataset allows for an interesting cross-validation approach that captures the multi-site nature of it. Our data was divided into 17 cross-validation sets. In each one, one site was used as the test set and the other 16 sites were used as the training set.

### 2.4 Model Training and Evaluation

In this study, we used a multilayer perceptron with two hidden layers. The model was implemented with Keras and using the Tensorflow 1.13 backend [37]. This model was chosen due to its ability to construct data driven features before using them for the final classification task. Both layers consisted of 512 rectified linear neurons. Due to the limited number of samples, the model was heavily regularized. The first hidden layer is subjected to L1 regularization in order to force the network to have some feature selection capabilities. A dropout layer is then applied before the second hidden layer. L2 regularization was used in the second hidden layer. Finally, a dropout layer was included before the output layer. The output layer is a single neuron which is activated by a sigmoid function.

The model training process is as follows. First, each feature was standardized by removing the mean and scaling to unit variance. Then the training data was ran through the neural network. The model used binary cross-entropy as its loss function and an Adam optimizer with *β*_l_ = 0.99, *β*_2_ = 0.01, and an initial learning rate of *l* = 0.0002. The learning rate was decayed by a factor of 0.5 base on the validation loss calculated on a random 10% of the training data withheld during the process. The model parameters were tuned on only one of the left out sites (PITT) to ensure low information leak. The trained model was then evaluated on the test set. We repeated the above steps 5 times for each LOSO fold and report the average accuracy, sensitivity, specificity and AUC score over the 5 repetitions. It is worth noting that the healthy subjects were given a label of 1 and the ASD subjects were given label 0. Thus, sensitivity should be interpreted as the percentage of HCs correctly identified. Likewise, specificity is the percentage of ASDs correctly identified.

### 2.5 Statistical Evaluation

We performed two statistical analyses to compare model results for each left out site between different pipelines. First, we compared the 5 trials performance of the best model for each pipeline for each site against each other using a Welch’s t-test [38]. We also compared model performance in a threshold-wise manner, comparing each threshold results for each pipeline against the same threshold results in another pipeline using the same test. The significance level is set to p<0.05.

## 3 Results

In order to manage the different thresholds, we report the performance measures of the best performing threshold according to AUC score for each site. This applies to all following paragraphs unless explicitly specified. The effects of this choice are discussed further in the discussion.

Our results (Tables 2-A to C) show that, on average, the negative correlation pipeline can achieve higher accuracies and AUC score than the other two pipelines. This is mostly due to a rise in specificity. The average sensitivity of the model remains unchanged over the pipelines while being slightly lower for the absolute value pipeline. Specifically, the model was able to achieve a higher specificity, increasing its ability to correctly identify ASDs. This is interesting because it suggests different feature extraction pipelines will allow some flexibility between interchanging sensitivity and specificity.

**Table 2-A.**
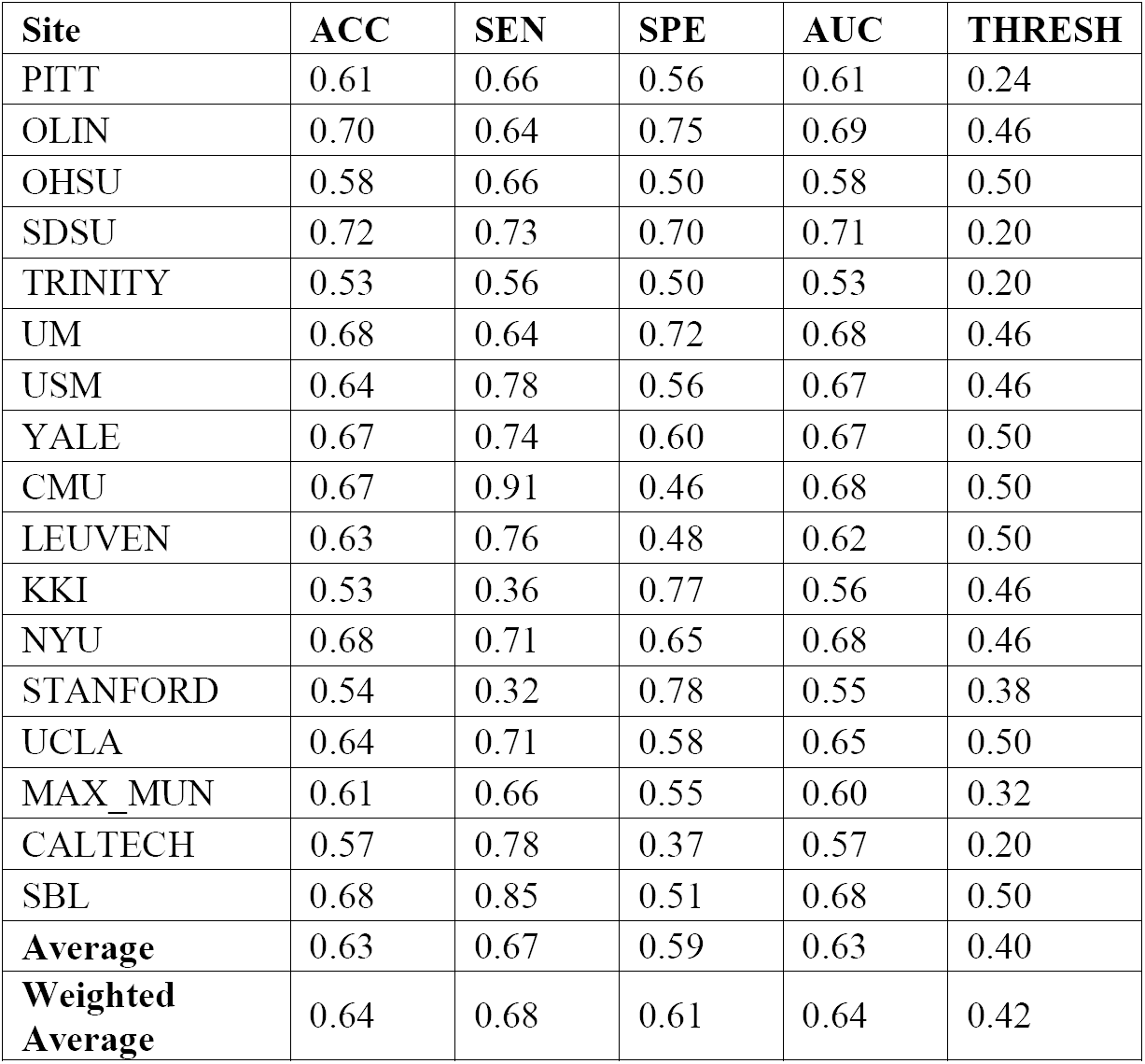
Results using the unchanged correlation matrix. ACC: ACURACY, SEN: SENSITIVITY, SPE: SPECIFICITY, AUC: Area under curve score, THRESH: Graph density threshold for the reported metrics

| Site | ACC | SEN | SPE | AUC | THRESH |
| --- | --- | --- | --- | --- | --- |
| PITT | 0.61 | 0.66 | 0.56 | 0.61 | 0.24 |
| OLIN | 0.70 | 0.64 | 0.75 | 0.69 | 0.46 |
| OHSU | 0.58 | 0.66 | 0.50 | 0.58 | 0.50 |
| SDSU | 0.72 | 0.73 | 0.70 | 0.71 | 0.20 |
| TRINITY | 0.53 | 0.56 | 0.50 | 0.53 | 0.20 |
| UM | 0.68 | 0.64 | 0.72 | 0.68 | 0.46 |
| USM | 0.64 | 0.78 | 0.56 | 0.67 | 0.46 |
| YALE | 0.67 | 0.74 | 0.60 | 0.67 | 0.50 |
| CMU | 0.67 | 0.91 | 0.46 | 0.68 | 0.50 |
| LEUVEN | 0.63 | 0.76 | 0.48 | 0.62 | 0.50 |
| KKI | 0.53 | 0.36 | 0.77 | 0.56 | 0.46 |
| NYU | 0.68 | 0.71 | 0.65 | 0.68 | 0.46 |
| STANFORD | 0.54 | 0.32 | 0.78 | 0.55 | 0.38 |
| UCLA | 0.64 | 0.71 | 0.58 | 0.65 | 0.50 |
| MAX_MUN | 0.61 | 0.66 | 0.55 | 0.60 | 0.32 |
| CALTECH | 0.57 | 0.78 | 0.37 | 0.57 | 0.20 |
| SBL | 0.68 | 0.85 | 0.51 | 0.68 | 0.50 |
| **Average** | 0.63 | 0.67 | 0.59 | 0.63 | 0.40 |
| **Weighted Average** | 0.64 | 0.68 | 0.61 | 0.64 | 0.42 |

**Table 2-B.** Results using the absolute value of the correlation matrix

|  | ACC | SEN | SPE | AUC | THRESH |
| --- | --- | --- | --- | --- | --- |
| PITT | 0.66 | 0.68 | 0.64 | 0.66 | 0.46 |
| OLIN | 0.68 | 0.55 | 0.79 | 0.67 | 0.46 |
| OHSU | 0.53 | 0.47 | 0.60 | 0.54 | 0.22 |
| SDSU | 0.66 | 0.69 | 0.61 | 0.65 | 0.20 |
| TRINITY | 0.58 | 0.59 | 0.56 | 0.58 | 0.48 |
| UM | 0.68 | 0.63 | 0.75 | 0.69 | 0.28 |
| USM | 0.65 | 0.75 | 0.60 | 0.67 | 0.44 |
| YALE | 0.65 | 0.81 | 0.49 | 0.65 | 0.24 |
| CMU | 0.65 | 0.75 | 0.56 | 0.66 | 0.30 |
| LEUVEN | 0.58 | 0.76 | 0.36 | 0.56 | 0.38 |
| KKI | 0.59 | 0.47 | 0.75 | 0.61 | 0.44 |
| NYU | 0.67 | 0.69 | 0.65 | 0.67 | 0.24 |
| STANFORD | 0.58 | 0.48 | 0.68 | 0.58 | 0.46 |
| UCLA | 0.68 | 0.69 | 0.66 | 0.68 | 0.44 |
| MAX_MUN | 0.61 | 0.69 | 0.52 | 0.60 | 0.30 |
| CALTECH | 0.61 | 0.66 | 0.56 | 0.61 | 0.24 |
| SBL | 0.59 | 0.84 | 0.35 | 0.59 | 0.20 |
| Average | 0.63 | 0.66 | 0.60 | 0.63 | 0.34 |
| Weighted<br>Average | 0.64 | 0.67 | 0.62 | 0.64 | 0.34 |

**Table 2-C.** Results using the negative correlation matrix

|  | ACC | SEN | SPE | AUC | THRESH |
| --- | --- | --- | --- | --- | --- |
| PITT | 0.66 | 0.79 | 0.54 | 0.67 | 0.28 |
| OLIN | 0.68 | 0.64 | 0.71 | 0.67 | 0.38 |
| OHSU | 0.48 | 0.47 | 0.50 | 0.49 | 0.46 |
| SDSU | 0.73 | 0.65 | 0.86 | 0.76 | 0.30 |
| TRINITY | 0.60 | 0.62 | 0.57 | 0.59 | 0.28 |
| UM | 0.69 | 0.62 | 0.77 | 0.70 | 0.44 |
| USM | 0.70 | 0.82 | 0.63 | 0.73 | 0.42 |
| YALE | 0.68 | 0.66 | 0.69 | 0.68 | 0.38 |
| CMU | 0.70 | 0.85 | 0.56 | 0.70 | 0.28 |
| LEUVEN | 0.65 | 0.80 | 0.48 | 0.64 | 0.46 |
| KKI | 0.61 | 0.46 | 0.81 | 0.64 | 0.20 |
| NYU | 0.65 | 0.66 | 0.63 | 0.65 | 0.46 |
| STANFORD | 0.58 | 0.39 | 0.78 | 0.58 | 0.44 |
| UCLA | 0.71 | 0.74 | 0.69 | 0.71 | 0.32 |
| MAX_MUN | 0.62 | 0.63 | 0.61 | 0.62 | 0.34 |
| CALTECH | 0.61 | 0.73 | 0.48 | 0.61 | 0.32 |
| SBL | 0.66 | 0.81 | 0.51 | 0.66 | 0.50 |
| Average | 0.65 | 0.67 | 0.64 | 0.65 | 0.37 |
| Weighted<br>Average | 0.66 | 0.67 | 0.65 | 0.66 | 0.38 |

The same results held when adding the PCA transformation of the original correlation matrix as input features. However, the overall performance of the model was improved. Suggesting that information from the original correlation matrix supplements the information available in only graph features (Supplementary tables 1-A to C). To test whether these observations were dependent on the choice of the atlas, we applied the same methodology on a different atlas When using the AAL parcellation, the negative pipeline performed the best as well. Thus, strengthening the hypothesis that this result holds across different parcellation atlases (Supplementary tables 2-A to C). Another variable in the dataset was gender. The negative correlation pipeline was still the best performing pipeline on average when only modelling the male subjects. Interestingly, overall model performance was lowered when the training and test data only included from male subjects. (Supplementary tables 3-A to C).

As evident in the performance tables, there is a high variability in model performance between sites. One possible explanation for this may be related to the age of the subjects in that site. To investigate this, we plotted the AUC scores of the 5 trials for each site against their age means. A quadratic regression was applied on this data and is shown in Figure 2. In the case of added PCA features, the sites at opposing ends of the age mean scale performed worse than the sites closer to the mean. This phenomenon was less pronounced in the case of using only graph features with it being almost non-existent in the case of the positive correlation pipeline.

**Figure 1.**
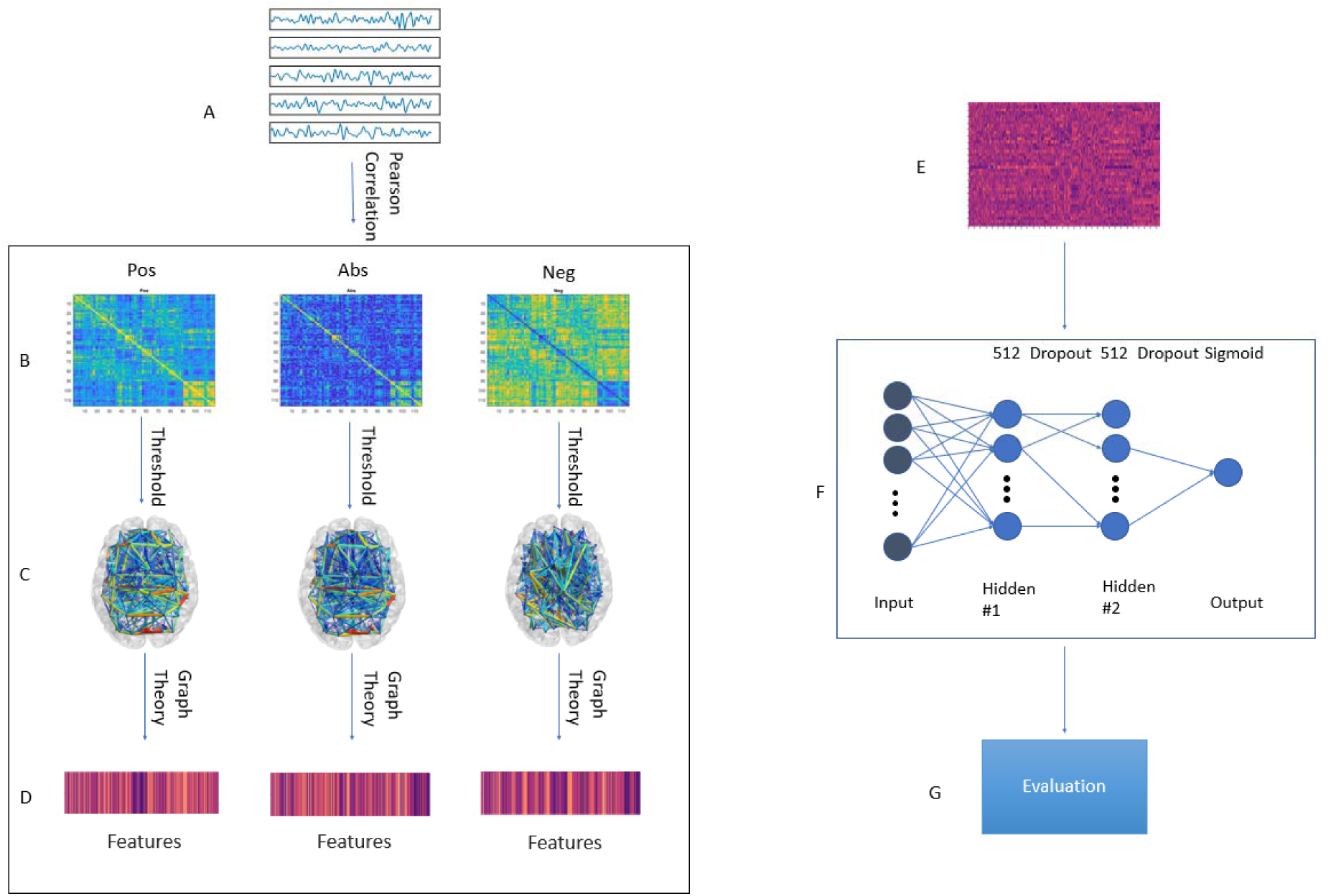
Graphical framework of the experiment. A. By averaging the BOLD activity in each ROI in the parcellation atlas, a time series is extracted representing brain activity in that region;Using Pearson’s correlation, a connectivity matrix is generated from the ROI time series quantifying the connectivity level between individual ROIs; The connectivity matrix is transformed using three pipelines: Pos. No change to the connectivity matrix, Neg: Multiplying the connectivity matrix by −1, Abs: calculating the absolute value of the matrix. Then, by treating the ROIs as graph nodes and the connectivity matrix as graph weights the brain network is expressed in graph form; A threshold is applied to keep only the 20-50% strongest connections in 2% increments; Graph theoretical analysis is applied to the resulting graph from to obtain a feature vector for each subject; E: Feature matrix for all subjects in the training fold; F. MLP architecture. First ReLU layer is l1-regularized and second ReLU layer is l2-regularized. A dropout of 0.7 was applied between the first and second ReLU as well as the second ReLU and output; G. The model is tested on a previously unseen test set from a different site in the ABIDE dataset. The evaluation metrics are: Accuracy, Sensitivity, Specificity, AUC score.

**Figure 2.**
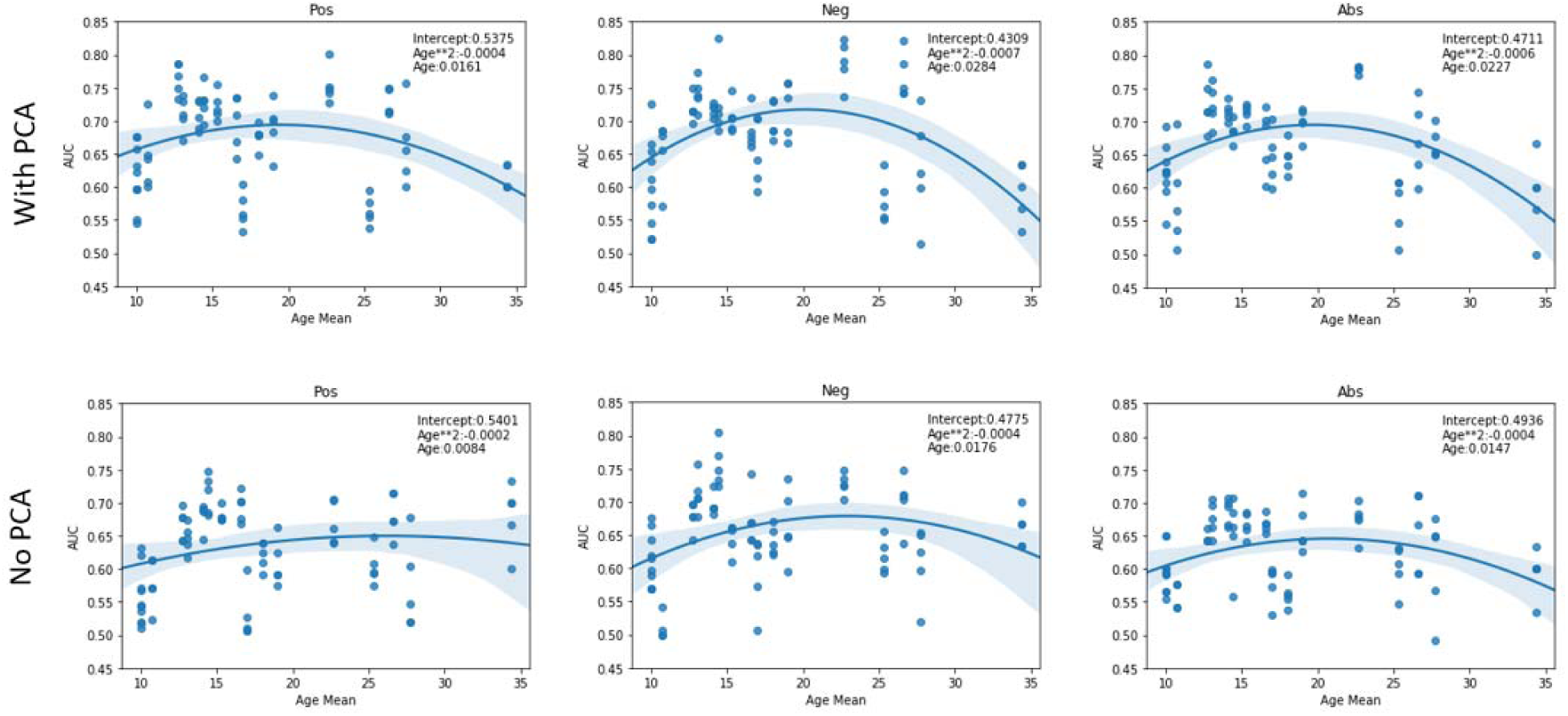
AUC scores vs Age. The figures show the quadratic regression line of AUC score of a LOSO fold based on mean age associated with the left-out site. Top row plots the results from using both PCA and graph features while bottom row shows result from using only graph features. The sites that have extreme age ranges (either lowest or highest) perform worse than those closer to the population median. An inverted U-shape is observed in all pipelines other than the Pos pipeline in the case of using only graph features. Abbreviations: Pos: Positive pipeline, Neg: negative pipeline, Abs, Absolute value pipeline

Supplementary tables Table 4-A and B show the p-values of between pipeline differences compared to the anti-correlation pipeline for the graph-features-only method. When comparing the best models, based on what threshold led to the highest AUC score, trained with only graph features for each site, the negative pipeline achieved a higher AUC than the other pipelines except in the following sites. OLIN, OHSU, NYU and SBL for the unchanged pipeline and OHSU and NYU for the absolute value pipeline. Of these sites, Only OHSU (p = 0.024) and NYU (p = 0.020) were significantly different (p<0.05) between the unchanged and negative pipelines. For accuracy, the same comparison holds between the negative and unchanged pipelines. OSHU (p = 0.014) and NYU (p = 0.014) results were again significantly different. The absolute value pipeline was able to achieve better accuracies than the negative pipeline for the following site: OLIN, OHSU, NYU. Likewise, the AAL atlas results did not show many statistical significances in the site accuracies between different pipelines.

As it may not be fair to compare different thresholds for each site, we also performed threshold matched comparisons for the graph-only craddock-200 atlas results. A Welch’s t-test was performed on the performance metrics of each pipeline pair for each site. Figure 3 shows these comparisons in a heatmap style plot. Interestingly, no site showed consistent difference in performance over the tested threshold range, suggesting that this value should be treated as a parameter than can affect the accuracy of machine learning predictions using graph theory and tuned for the task at hand.

**Figure 3.**
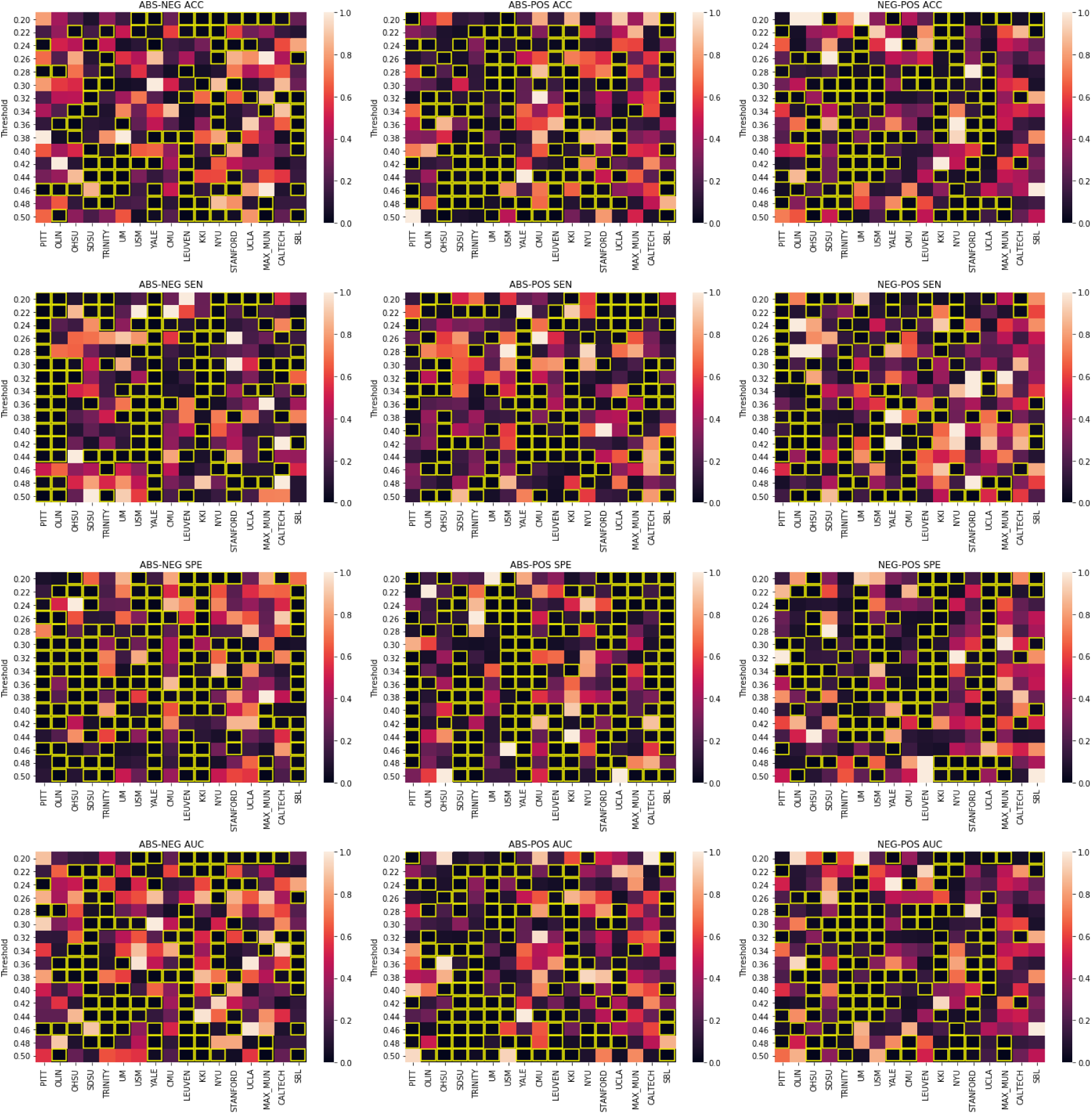
Welch’s t-test results, Figure columns represent the pipelines being compared and the rows represent the metric that is being compared. In each sub-figure, the title shows the pipelines being compared following by the metric. The X-axis represents different imaging sites and the Y-axis shows different thresholds. The heatmap values are p-values distributed based on the colormap to the right of each figure. Significant p-values(0.05) are highlighted in yellow

Although not the main goal of this study, our model was able to, on average, perform on-par with the deep learning model described in the Heinsfeld et al paper[23] when only using the graph features. The sites CALTECH, KKI, MAX_MUN, NYU, OHSU, STANFORD and TRINITY showcased lower accuracy in our model. However, when the PCA features were also used as input features, our model was able to achieve an average accuracy of 68% and a weighted average (based on the test set size) of 69%. The sites that still showcased lower accuracy were CALTECH, KKI, MAX_MUN, SBL and STANFORD.

## 4 Discussion

Our results consistently showed that using graph theoretical features from the negative pipeline increases the MLP model accuracy in distinguishing between HC and ASD. This suggests that anti-correlations in the brain network contain important information that aids our model in distinguishing between ASD and HC. One reason for the higher performance of the anti-correlation method could be that contrary to some previous studies [16][17], GSR introduced anti-correlations may in fact stem from real neurological basis and not spurious [38]. Unfortunately, as we do not know of any gold standards to test this hypothesis in real data, it should only be interpreted as a speculation made based on the results of this study.

Previous studies have concluded that the use of absolute value graph metrics in the presence of GSR may be compromised [37]. This was attributed to the fact that in the presence of GSR, the topologies of the anti-correlation and correlation matrix will be mixed when using an absolute value pipeline as the anti-correlations have comparable magnitude to the correlations. However, our model trained with the absolute value graph features was able to perform on-par with the positive correlation pipeline.

The small amount of significantly different within-site metrics between the three pipelines may be attributed to the way they were calculated. Each leave-one-site fold was run 5 times in order to offset any effects of random weight initialization and random training-validation set splits. Thus, increasing the number of repetitions may lead to higher levels of difference significance between the reported metrics. Another possible explanation is that there is enough information available in all pipelines for most of these sites.

In the case of added PCA, plotting the accuracies of the negative pipeline against number of participants did not provide any further insight into why some sites performed worst. Plotting the same against the age of the participants revealed that there may be a link between age and the ability of our model to make accurate predictions. Age has been previously shown to affect network properties of subject within the ABIDE dataset [39]. KKI and STANFORD are the two sites with the lowest average ASD and HC age. Of the 4 sites with the highest average ASD age, 3 of them, CALTECH, MAX_MUN and SBL showed lower accuracies. Interestingly, CMU, did not follow this behavior. However, the standard deviation of the age of CMU’s participants was lower than those of the other 3. The larger age SD may have hindered the algorithm’s ability to perform well for the sites with high average age.

Interestingly, the model trained only on the graph features was able to perform relatively well on SBL despite that site having the highest age mean and performed worst on OHSU, the site with the least number of subjects. While this could have happened purely by chance, it is an interesting avenue for further insights about the nature of our model. Furthermore, these models were more robust to the effects of age as shown in figure 2 suggesting age-related information is better captured in the graph theoretical features than the PCA transformation of the correlation matrix.

Another intriguing observation about our results is that some thresholds and pipelines were better suited for classifying specific sites. This suggest that an ensemble model, such as training different models on each of these combinations and assigning the most selected label for each subject between these models, may achieve better performance on this dataset.

In this study we utilized a leave-one-site out cross validation approach in order to capture the multi-site nature of the ABIDE I dataset. We believe this approach leads to a better estimate of how this model will perform in the real world as it allows for better generalization across previously unseen imaging sites and protocols.

Using graph theoretical measures to analyze and classify neurological and neurodevelopmental diseases is not a novel idea. However, the practice of thresholding the connectivity matrix in order to produce a binary graph introduces an artificial bias towards some of the correlations. While this bias can be avoided by using the absolute value of the correlation matrix, this may disregard the important information that the positive correlation and anti-correlation networks hold. We have shown that, in the presence of GSR, focusing on the anti-correlation network lead to better classification performance in the case of ASD vs HC. We also showed that by adding non-graph features to the anti-correlation graph features, classification metrics are improved suggesting that a different process may be needed to accurately capture the graph properties of both the positive and negative correlation networks.

One limitation in the present study may be the use of multiple threshold values in constructing the functional graph. Here we reported the results from the threshold resulting in the highest AUC for each site. A fairer approach may be to average the results of all thresholds for each site. This led to a significant decrease in model performance however the negative correlation pipeline still outperformed the other pipeline.

Another limitation of this study was the inclusion of all age ranges in our dataset. As different sites have different demographics, this may affect the power of our model. Furthermore, identifying ASD is most important in earlier stages of life. The ABIDE I dataset included only a small number of children under the age of 10 (146 subjects) spread across multiple sites and none below the age of 5. Thus, the results presented in this study may not generalize to studies on younger children.

The ABIDE I data exhibits numerous inherent variability due to its multi-site nature. Here we presented a deep learning model that was able to navigate the intricacies of this data and generalize over multiple sites by using graph theoretical features. Our model was able to perform on-par or better than previously reported deep learning models using the same number of subjects.

## 5 Acknowledgments

Part of the code for the preprocessing pipeline was reused from the open source code provided by the Heinsfeld et al. paper. This work could not be completed without the immense contribution of open sourcing the ABIDE datasets.

