## Supplementary Tables for "The Importance of Anti-Correlations in Graph Theory Based Classification of Autism Spectrum Disorder"

Supplementary Material

**Dosenbach Atlas, PCA + Graph features:**

Table 1-A. Results using the unchanged correlation matrix. ACC: ACCURACY, SEN: SENSITIVITY, SPE: SPECIFICITY, AUC: Area under curve score, THRESH: Graph density threshold for the reported metrics

| **SITE** | **ACC** | **SEN** | **SPE** | **AUC** | **THRESH** |
| --- | --- | --- | --- | --- | --- |
| **PITT** | 0.69 | 0.84 | 0.54 | 0.69 | 0.50 |
| **OLIN** | 0.70 | 0.68 | 0.72 | 0.70 | 0.46 |
| **OHSU** | 0.65 | 0.76 | 0.53 | 0.65 | 0.46 |
| **SDSU** | 0.74 | 0.80 | 0.66 | 0.73 | 0.20 |
| **TRINITY** | 0.56 | 0.50 | 0.63 | 0.57 | 0.22 |
| **UM** | 0.70 | 0.63 | 0.78 | 0.71 | 0.28 |
| **USM** | 0.75 | 0.76 | 0.75 | 0.75 | 0.50 |
| **YALE** | 0.76 | 0.88 | 0.65 | 0.76 | 0.20 |
| **CMU** | 0.72 | 0.97 | 0.49 | 0.73 | 0.50 |
| **LEUVEN** | 0.70 | 0.91 | 0.44 | 0.68 | 0.50 |
| **KKI** | 0.61 | 0.49 | 0.79 | 0.64 | 0.26 |
| **NYU** | 0.72 | 0.70 | 0.74 | 0.72 | 0.20 |
| **STANFORD** | 0.58 | 0.33 | 0.85 | 0.59 | 0.48 |
| **UCLA** | 0.71 | 0.73 | 0.69 | 0.71 | 0.50 |
| **MAX_MUN** | 0.57 | 0.57 | 0.56 | 0.56 | 0.20 |
| **CALTECH** | 0.66 | 0.78 | 0.55 | 0.66 | 0.22 |
| **SBL** | 0.61 | 0.85 | 0.37 | 0.61 | 0.20 |
| **Average** | 0.67 | 0.72 | 0.63 | 0.67 | 0.35 |
| **Weighted Average** | 0.69 | 0.71 | 0.66 | 0.68 | 0.33 |

Table 1-B. Results using the absolute value of the correlation matrix

| **SITE** | **ACC** | **SEN** | **SPE** | **AUC** | **THRESH** |
| --- | --- | --- | --- | --- | --- |
| **PITT** | 0.71 | 0.87 | 0.57 | 0.72 | 0.44 |
| **OLIN** | 0.69 | 0.65 | 0.72 | 0.68 | 0.38 |
| **OHSU** | 0.66 | 0.74 | 0.57 | 0.65 | 0.46 |
| **SDSU** | 0.73 | 0.73 | 0.73 | 0.73 | 0.22 |
| **TRINITY** | 0.65 | 0.66 | 0.65 | 0.65 | 0.24 |
| **UM** | 0.71 | 0.65 | 0.78 | 0.72 | 0.44 |
| **USM** | 0.78 | 0.82 | 0.75 | 0.79 | 0.40 |
| **YALE** | 0.72 | 0.71 | 0.72 | 0.72 | 0.44 |
| **CMU** | 0.76 | 0.91 | 0.63 | 0.77 | 0.28 |
| **LEUVEN** | 0.71 | 0.88 | 0.52 | 0.70 | 0.30 |
| **KKI** | 0.62 | 0.41 | 0.91 | 0.66 | 0.44 |
| **NYU** | 0.71 | 0.71 | 0.70 | 0.71 | 0.44 |
| **STANFORD** | 0.54 | 0.24 | 0.86 | 0.55 | 0.32 |
| **UCLA** | 0.74 | 0.76 | 0.72 | 0.74 | 0.32 |
| **MAX_MUN** | 0.58 | 0.64 | 0.53 | 0.58 | 0.36 |
| **CALTECH** | 0.63 | 0.69 | 0.57 | 0.63 | 0.32 |
| **SBL** | 0.59 | 0.71 | 0.48 | 0.59 | 0.26 |
| **Average** | 0.68 | 0.69 | 0.67 | 0.68 | 0.36 |
| **Weighted Average** | 0.69 | 0.7 | 0.69 | 0.69 | 0.38 |

Table 1-C. Results using the negative correlation matrix

| **SITE** | **ACC** | **SEN** | **SPE** | **AUC** | **THRESH** |
| --- | --- | --- | --- | --- | --- |
| **PITT** | 0.70 | 0.76 | 0.63 | 0.70 | 0.38 |
| **OLIN** | 0.68 | 0.59 | 0.76 | 0.67 | 0.44 |
| **OHSU** | 0.58 | 0.61 | 0.55 | 0.58 | 0.28 |
| **SDSU** | 0.69 | 0.73 | 0.64 | 0.69 | 0.50 |
| **TRINITY** | 0.64 | 0.59 | 0.70 | 0.65 | 0.48 |
| **UM** | 0.71 | 0.61 | 0.82 | 0.71 | 0.34 |
| **USM** | 0.77 | 0.80 | 0.76 | 0.78 | 0.22 |
| **YALE** | 0.73 | 0.83 | 0.63 | 0.73 | 0.44 |
| **CMU** | 0.67 | 0.80 | 0.54 | 0.67 | 0.40 |
| **LEUVEN** | 0.66 | 0.88 | 0.41 | 0.65 | 0.46 |
| **KKI** | 0.62 | 0.48 | 0.81 | 0.64 | 0.36 |
| **NYU** | 0.72 | 0.74 | 0.69 | 0.71 | 0.34 |
| **STANFORD** | 0.59 | 0.29 | 0.92 | 0.60 | 0.26 |
| **UCLA** | 0.72 | 0.72 | 0.73 | 0.72 | 0.44 |
| **MAX_MUN** | 0.58 | 0.69 | 0.46 | 0.57 | 0.30 |
| **CALTECH** | 0.66 | 0.76 | 0.58 | 0.67 | 0.24 |
| **SBL** | 0.59 | 0.85 | 0.32 | 0.59 | 0.50 |
| **Average** | 0.67 | 0.69 | 0.64 | 0.67 | 0.38 |
| **Weighted Average** | 0.68 | 0.7 | 0.67 | 0.69 | 0.37 |

**AAL Atlas, Graph features:**

Table 2-A. Results using the unchanged correlation matrix. ACC: ACCURACY, SEN: SENSITIVITY, SPE: SPECIFICITY, AUC: Area under curve score, THRESH: Graph density threshold for the reported metrics

| **SITE** | **ACC** | **SEN** | **SPE** | **AUC** | **THRESH** |
| --- | --- | --- | --- | --- | --- |
| **PITT** | 0.55 | 0.61 | 0.50 | 0.56 | 0.28 |
| **OLIN** | 0.73 | 0.69 | 0.76 | 0.73 | 0.24 |
| **OHSU** | 0.58 | 0.51 | 0.65 | 0.58 | 0.22 |
| **SDSU** | 0.74 | 0.74 | 0.74 | 0.74 | 0.34 |
| **TRINITY** | 0.60 | 0.66 | 0.55 | 0.60 | 0.34 |
| **UM** | 0.61 | 0.54 | 0.69 | 0.61 | 0.20 |
| **USM** | 0.63 | 0.71 | 0.58 | 0.65 | 0.48 |
| **YALE** | 0.72 | 0.73 | 0.71 | 0.72 | 0.36 |
| **CMU** | 0.71 | 0.80 | 0.63 | 0.71 | 0.38 |
| **LEUVEN** | 0.55 | 0.74 | 0.32 | 0.53 | 0.34 |
| **KKI** | 0.54 | 0.53 | 0.56 | 0.54 | 0.38 |
| **NYU** | 0.62 | 0.64 | 0.59 | 0.61 | 0.50 |
| **STANFORD** | 0.57 | 0.43 | 0.72 | 0.57 | 0.24 |
| **UCLA** | 0.63 | 0.75 | 0.53 | 0.64 | 0.40 |
| **MAX_MUN** | 0.58 | 0.70 | 0.45 | 0.58 | 0.46 |
| **CALTECH** | 0.54 | 0.58 | 0.51 | 0.54 | 0.30 |
| **SBL** | 0.75 | 0.71 | 0.79 | 0.75 | 0.22 |
| **Average** | 0.63 | 0.65 | 0.60 | 0.63 | 0.33 |
| **Weighted Average** | 0.62 | 0.65 | 0.6 | 0.62 | 0.36 |

Table 2-B. Results using the absolute value of the correlation matrix

| **SITE** | **ACC** | **SEN** | **SPE** | **AUC** | **THRESH** |
| --- | --- | --- | --- | --- | --- |
| **PITT** | 0.63 | 0.81 | 0.45 | 0.63 | 0.34 |
| **OLIN** | 0.74 | 0.75 | 0.73 | 0.74 | 0.32 |
| **OHSU** | 0.63 | 0.59 | 0.68 | 0.63 | 0.24 |
| **SDSU** | 0.66 | 0.64 | 0.70 | 0.67 | 0.22 |
| **TRINITY** | 0.62 | 0.69 | 0.54 | 0.61 | 0.34 |
| **UM** | 0.64 | 0.53 | 0.75 | 0.64 | 0.28 |
| **USM** | 0.69 | 0.86 | 0.60 | 0.73 | 0.40 |
| **YALE** | 0.74 | 0.71 | 0.77 | 0.74 | 0.30 |
| **CMU** | 0.79 | 0.83 | 0.74 | 0.79 | 0.50 |
| **LEUVEN** | 0.63 | 0.82 | 0.41 | 0.62 | 0.22 |
| **KKI** | 0.58 | 0.54 | 0.64 | 0.59 | 0.30 |
| **NYU** | 0.62 | 0.53 | 0.74 | 0.64 | 0.38 |
| **STANFORD** | 0.54 | 0.38 | 0.72 | 0.55 | 0.32 |
| **UCLA** | 0.64 | 0.74 | 0.56 | 0.65 | 0.22 |
| **MAX_MUN** | 0.63 | 0.65 | 0.61 | 0.63 | 0.24 |
| **CALTECH** | 0.58 | 0.81 | 0.36 | 0.58 | 0.34 |
| **SBL** | 0.71 | 0.68 | 0.73 | 0.71 | 0.30 |
| **Average** | 0.65 | 0.68 | 0.63 | 0.66 | 0.31 |
| **Weighted Average** | 0.64 | 0.66 | 0.64 | 0.65 | 0.31 |

Table 2-C. Results using the negative correlation matrix

| **SITE** | **ACC** | **SEN** | **SPE** | **AUC** | **THRESH** |
| --- | --- | --- | --- | --- | --- |
| **PITT** | 0.57 | 0.61 | 0.53 | 0.57 | 0.50 |
| **OLIN** | 0.69 | 0.65 | 0.72 | 0.68 | 0.28 |
| **OHSU** | 0.65 | 0.51 | 0.80 | 0.66 | 0.32 |
| **SDSU** | 0.66 | 0.55 | 0.83 | 0.69 | 0.42 |
| **TRINITY** | 0.62 | 0.70 | 0.53 | 0.61 | 0.44 |
| **UM** | 0.63 | 0.65 | 0.59 | 0.62 | 0.36 |
| **USM** | 0.64 | 0.83 | 0.53 | 0.68 | 0.20 |
| **YALE** | 0.71 | 0.74 | 0.67 | 0.71 | 0.20 |
| **CMU** | 0.74 | 0.86 | 0.63 | 0.75 | 0.48 |
| **LEUVEN** | 0.60 | 0.76 | 0.41 | 0.59 | 0.20 |
| **KKI** | 0.59 | 0.54 | 0.66 | 0.60 | 0.24 |
| **NYU** | 0.58 | 0.61 | 0.55 | 0.58 | 0.36 |
| **STANFORD** | 0.67 | 0.52 | 0.82 | 0.67 | 0.34 |
| **UCLA** | 0.65 | 0.70 | 0.61 | 0.65 | 0.36 |
| **MAX_MUN** | 0.57 | 0.57 | 0.58 | 0.57 | 0.22 |
| **CALTECH** | 0.59 | 0.52 | 0.65 | 0.59 | 0.44 |
| **SBL** | 0.67 | 0.64 | 0.71 | 0.67 | 0.24 |
| **Average** | 0.64 | 0.65 | 0.64 | 0.64 | 0.33 |
| **Weighted Average** | 0.62 | 0.65 | 0.6 | 0.63 | 0.33 |

**Dosenbach, Graph features, Only Males:**

Table 3-A. Results using the unchanged correlation matrix. ACC: ACCURACY, SEN: SENSITIVITY, SPE: SPECIFICITY, AUC: Area under curve score, THRESH: Graph density threshold for the reported metrics

| **SITE** | **ACC** | **SEN** | **SPE** | **AUC** | **THRESH** |
| --- | --- | --- | --- | --- | --- |
| **PITT** | 0.63 | 0.67 | 0.60 | 0.63 | 0.30 |
| **OLIN** | 0.72 | 0.58 | 0.83 | 0.70 | 0.48 |
| **OHSU** | 0.57 | 0.60 | 0.53 | 0.57 | 0.50 |
| **SDSU** | 0.71 | 0.59 | 0.86 | 0.72 | 0.22 |
| **TRINITY** | 0.51 | 0.56 | 0.46 | 0.51 | 0.20 |
| **UM** | 0.65 | 0.55 | 0.74 | 0.65 | 0.50 |
| **USM** | 0.62 | 0.70 | 0.57 | 0.64 | 0.46 |
| **YALE** | 0.70 | 0.70 | 0.69 | 0.70 | 0.50 |
| **CMU** | 0.65 | 0.78 | 0.53 | 0.65 | 0.20 |
| **LEUVEN** | 0.63 | 0.81 | 0.43 | 0.62 | 0.48 |
| **KKI** | 0.51 | 0.23 | 0.85 | 0.54 | 0.50 |
| **NYU** | 0.67 | 0.63 | 0.72 | 0.68 | 0.20 |
| **STANFORD** | 0.56 | 0.28 | 0.87 | 0.57 | 0.44 |
| **UCLA** | 0.65 | 0.64 | 0.66 | 0.65 | 0.46 |
| **MAX_MUN** | 0.64 | 0.61 | 0.68 | 0.65 | 0.34 |
| **CALTECH** | 0.59 | 0.64 | 0.53 | 0.59 | 0.34 |
| **SBL** | 0.69 | 0.80 | 0.57 | 0.69 | 0.26 |
| **Average** | 0.63 | 0.61 | 0.65 | 0.63 | 0.38 |
| **Weighted Average** | 0.64 | 0.61 | 0.67 | 0.64 | 0.38 |

Table 3-B. Results using the absolute value of the correlation matrix

| **SITE** | **ACC** | **SEN** | **SPE** | **AUC** | **THRESH** |
| --- | --- | --- | --- | --- | --- |
| **PITT** | 0.65 | 0.70 | 0.61 | 0.66 | 0.22 |
| **OLIN** | 0.68 | 0.57 | 0.76 | 0.67 | 0.44 |
| **OHSU** | 0.46 | 0.40 | 0.53 | 0.47 | 0.46 |
| **SDSU** | 0.68 | 0.55 | 0.83 | 0.69 | 0.30 |
| **TRINITY** | 0.56 | 0.63 | 0.48 | 0.56 | 0.24 |
| **UM** | 0.70 | 0.64 | 0.75 | 0.70 | 0.40 |
| **USM** | 0.64 | 0.82 | 0.53 | 0.68 | 0.44 |
| **YALE** | 0.67 | 0.62 | 0.71 | 0.67 | 0.34 |
| **CMU** | 0.72 | 0.86 | 0.60 | 0.73 | 0.28 |
| **LEUVEN** | 0.65 | 0.77 | 0.53 | 0.65 | 0.38 |
| **KKI** | 0.59 | 0.31 | 0.95 | 0.63 | 0.50 |
| **NYU** | 0.62 | 0.66 | 0.58 | 0.62 | 0.44 |
| **STANFORD** | 0.56 | 0.28 | 0.87 | 0.57 | 0.44 |
| **UCLA** | 0.65 | 0.62 | 0.68 | 0.65 | 0.20 |
| **MAX_MUN** | 0.64 | 0.64 | 0.63 | 0.64 | 0.26 |
| **CALTECH** | 0.67 | 0.79 | 0.56 | 0.67 | 0.32 |
| **SBL** | 0.66 | 0.84 | 0.48 | 0.66 | 0.32 |
| **Average** | 0.64 | 0.63 | 0.65 | 0.64 | 0.35 |
| **Weighted Average** | 0.64 | 0.64 | 0.65 | 0.65 | 0.36 |

Table 3-C. Results using the negative correlation matrix

| **SITE** | **ACC** | **SEN** | **SPE** | **AUC** | **THRESH** |
| --- | --- | --- | --- | --- | --- |
| **PITT** | 0.68 | 0.64 | 0.71 | 0.68 | 0.42 |
| **OLIN** | 0.61 | 0.42 | 0.78 | 0.60 | 0.50 |
| **OHSU** | 0.55 | 0.51 | 0.60 | 0.56 | 0.22 |
| **SDSU** | 0.66 | 0.56 | 0.78 | 0.67 | 0.44 |
| **TRINITY** | 0.61 | 0.70 | 0.51 | 0.60 | 0.22 |
| **UM** | 0.67 | 0.64 | 0.71 | 0.67 | 0.40 |
| **USM** | 0.63 | 0.77 | 0.56 | 0.66 | 0.44 |
| **YALE** | 0.67 | 0.79 | 0.54 | 0.67 | 0.22 |
| **CMU** | 0.60 | 0.76 | 0.45 | 0.61 | 0.30 |
| **LEUVEN** | 0.61 | 0.76 | 0.45 | 0.61 | 0.32 |
| **KKI** | 0.63 | 0.54 | 0.75 | 0.65 | 0.32 |
| **NYU** | 0.66 | 0.62 | 0.70 | 0.66 | 0.26 |
| **STANFORD** | 0.57 | 0.26 | 0.89 | 0.58 | 0.38 |
| **UCLA** | 0.63 | 0.68 | 0.59 | 0.64 | 0.34 |
| **MAX_MUN** | 0.62 | 0.59 | 0.65 | 0.62 | 0.30 |
| **CALTECH** | 0.63 | 0.76 | 0.52 | 0.64 | 0.50 |
| **SBL** | 0.63 | 0.84 | 0.43 | 0.63 | 0.50 |
| **Average** | 0.63 | 0.64 | 0.63 | 0.63 | 0.36 |
| **Weighted Average** | 0.64 | 0.65 | 0.64 | 0.64 | 0.36 |

**P-values of statistical test 1 as explained in the main body:**

Table 3-A. p-values between the results from the anticorrelation pipeline and absolute value pipelines. Significant values are shown in green (p<0.05)

|  | **ACC** | **SEN** | **SPE** | **AUC** |
| --- | --- | --- | --- | --- |
| **PITT** | 1 | 0.00704 | 0.049088 | 0.905637 |
| **OLIN** | 0.745423 | 0.090018 | 0.019257 | 0.821816 |
| **OHSU** | 0.264679 | 1 | 0.094141 | 0.214987 |
| **SDSU** | 0.063638 | 0.433303 | 0.00004 | 0.01156 |
| **TRINITY** | 0.555322 | 0.372966 | 0.831455 | 0.573903 |
| **UM** | 0.501903 | 0.869464 | 0.391634 | 0.444736 |
| **USM** | 0.024404 | 0.033199 | 0.29734 | 0.006976 |
| **YALE** | 0.026911 | 0.001852 | 0.000223 | 0.026911 |
| **CMU** | 0.193015 | 0.300643 | 1 | 0.192664 |
| **LEUVEN** | 0.00028 | 0.242119 | 0.013745 | 0.000275 |
| **KKI** | 0.420509 | 0.849664 | 0.143648 | 0.304904 |
| **NYU** | 0.099042 | 0.149582 | 0.627434 | 0.130409 |
| **STANFORD** | 1 | 0.017178 | 0.041921 | 0.854016 |
| **UCLA** | 0.077851 | 0.070563 | 0.23401 | 0.070927 |
| **MAX_MUN** | 0.545763 | 0.03776 | 0.03403 | 0.390427 |
| **CALTECH** | 1 | 0.077971 | 0.375713 | 0.962579 |
| **SBL** | 0.012924 | 0.242876 | 0.00619 | 0.012924 |

Table 3-B. p-values between the results from the anticorrelation pipeline and positive pipelines. Significant values are shown in green (p<0.05)

|  | **ACC** | **SEN** | **SPE** | **AUC** |
| --- | --- | --- | --- | --- |
| **PITT** | 0.111491 | 0.001875 | 0.758752 | 0.095194 |
| **OLIN** | 0.266627 | 1 | 0.195389 | 0.34826 |
| **OHSU** | 0.014456 | 0.000045 | 1 | 0.024348 |
| **SDSU** | 0.321425 | 0.137486 | 0.060976 | 0.062476 |
| **TRINITY** | 0.065748 | 0.054074 | 0.175743 | 0.069965 |
| **UM** | 0.309386 | 0.389027 | 0.038197 | 0.243787 |
| **USM** | 0.0319 | 0.0203 | 0.097081 | 0.014103 |
| **YALE** | 0.481453 | 0.038982 | 0.020157 | 0.481453 |
| **CMU** | 0.360802 | 0.308386 | 0.103791 | 0.42871 |
| **LEUVEN** | 0.127988 | 0.074681 | 0.698401 | 0.144683 |
| **KKI** | 0.019064 | 0.028193 | 0.332427 | 0.023005 |
| **NYU** | 0.014188 | 0.015801 | 0.447723 | 0.020558 |
| **STANFORD** | 0.143854 | 0.083119 | 1 | 0.151991 |
| **UCLA** | 0.002746 | 0.423281 | 0.001345 | 0.003386 |
| **MAX_MUN** | 0.521641 | 0.37543 | 0.060718 | 0.403319 |
| **CALTECH** | 0.369812 | 0.412668 | 0.193263 | 0.388424 |
| **SBL** | 0.466957 | 0.206151 | 1 | 0.466957 |
